# Noninvasive Assessment of Temporal Dynamics in Sympathetic and Parasympathetic Baroreflex Responses

**DOI:** 10.1101/2024.10.11.617927

**Authors:** Heberto Suarez-Roca, Negmeldeen Mamoun, Joseph P Mathew, Andrey V Bortsov

## Abstract

**Background:** The baroreflex system is crucial for cardiovascular regulation and autonomic homeostasis. A comprehensive assessment requires understanding the simultaneous temporal dynamics of its multiple functional branches, which traditional methods often overlook.

**Objective:** To develop and validate a noninvasive method for simultaneously assessing the temporal dynamics of sympathetic and parasympathetic baroreflexes using pulse contour analysis and the sequence method.

**Methods:** Beat-to-beat blood pressure and ECG recordings were analyzed from 55 preoperative cardiothoracic surgery patients in the supine position and 21 subjects from the EUROBAVAR dataset in both supine and standing positions. Systolic arterial pressure (SAP), interbeat interval (IBI), cardiac output (CO), myocardial contraction (dP/dt_max_), and systemic vascular resistance (SVR) were estimated using pulse contour analysis. Baroreflex sensitivity (BRS) was calculated via the sequence method and correlated with hemodynamic and heart rate variability (HRV) parameters.

**Results:** Parasympathetic BRS for IBI was correlated with the root mean square of successive differences of ECG RR intervals (RMSSD-HRV) at 0-beat delay. Sympathetic BRS for SVR strongly correlated with SVR, CO, and RMSSD-HRV, particularly at 3-beat delay, and was uniquely associated with SAP at 1-beat delay. Sympathetic BRS for dP/dt_max_ correlated with dP/dt_max_ at 1-beat delay. In contrast, BRS for CO correlated with CO and SVR at 0- and 3-beat delays. Postural changes mainly affected parasympathetically-mediated BRS for IBI and, to a lesser extent, the sympathetic vascular and myocardial branches.

**Conclusions:** This method effectively captures multiple baroreflex responses and their temporal dynamics, revealing distinct autonomic mechanisms and the impact of postural changes. Further validation is warranted.

## 1. INTRODUCTION

The complex interplay between baroreflexes, the autonomic nervous system, and hemodynamic regulation is crucial for maintaining cardiovascular control and autonomic homeostasis (Suarez-Roca et al., 2024). The arterial baroreflex system operates through a dynamic balance through two autonomically controlled closed loops: the parasympathetic baroreflex, which controls heart rate, and the sympathetic baroreflexes, which influence myocardial contraction and vascular tone (Båth et al., 1981; Ursino, 1999). Disruptions in this balance are evident in conditions such as heart failure (Dyavanapalli, 2020), diabetes mellitus (Sakamoto et al., 2019), obstructive sleep apnea (Labarca et al., 2020), chronic kidney disease (Salman, 2015; Soomro et al., 2021), and neurodegenerative disorders (Huang et al., 2022; Sabino-Carvalho et al., 2021), increasing the risk of acute kidney dysfunction (Ranucci et al., 2017) and cardiovascular events (La Rovere et al., 1998; La Rovere et al., 2001), especially in elderly populations (Fu et al., 2019).

The parasympathetic baroreflex is typically evaluated by analyzing changes in heart rate in response to blood pressure variations using time and frequency domain methods (Parati et al., 2000a). While effective in detecting dysfunction in cases like hypertension (Freitas et al., 2017), this method may not fully capture the severity of autonomic impairment, particularly in conditions like Parkinson’s disease (Huang et al., 2022). Evaluating the sympathetic baroreflex could provide additional insights, though it typically requires invasive procedures, such as recording muscle sympathetic nerve activity (Parati et al., 2012), which includes only vasoconstrictor nerve fibers and does not correlate with mean blood pressure in young adults (Wallin et al., 2007) or the sympathetic nerve activity of other areas, such as skin (Grassi et al., 1998).

Noninvasive methods, such as cross-spectral analysis between beat-to-beat blood pressure and stroke volume from impedance cardiography or the Modelflow method, provide alternatives for assessing sympathetic baroreflexes (Vaschillo et al., 2012; Yasumasu et al., 2004). Stroke volume, derived from pulse contour analysis, can be used to calculate systemic vascular resistance (SVR), and both can be cross-correlated with blood pressure oscillations (Borgers et al., 2014). However, these methods are affected by factors like heart rate, severe ectopic activity (La Rovere et al., 2008), low coherence (e.g., chronic heart failure), and non-baroreflex influences (Ondrusova et al., 2017; Pinna et al., 2002). The sequence method, analyzing the systolic pre-ejection period from thoracic impedance cardiography, is less affected by heart rate and preload factors (Reyes Del Paso et al., 2017) but often results in few valid baroreflex sequences (Reyes Del Paso et al., 2021). Furthermore, baroreflex responses of the sinus atrial node, myocardium, and vascular tone exhibit inherent and distinctive latencies relative to blood pressure changes (Vaschillo et al., 2012). These latencies depend on anatomical factors and neural mechanisms and are altered in conditions like resistant hypertension (Freitas et al., 2017). However, it is unclear how these latencies are associated with the ongoing levels of hemodynamics parameters or whether they change in response to orthostatic challenges.

This study aimed to develop a noninvasive methodology to capture the temporal dynamics of multiple baroreflex responses and validate its effectiveness. We assessed sympathetic and parasympathetic baroreflexes by combining noninvasive estimation of real-time, beat-to-beat hemodynamic parameters with pulse contour analysis (Saugel et al., 2021) and the sequence method for estimating baroreflex sensitivity (BRS) (Bertinieri et al., 1985; Parati et al., 2000a). We quantified changes in SVR, myocardial contraction, cardiac output, and interbeat interval in response to changes in SAP. We analyzed these responses at various beat delays following the pressure pulse to characterize the baroreflex’s temporal dynamics and autonomic nature. We validated these parameters by correlating them with key hemodynamic parameters and assessing the impact of positional changes. We hypothesized that this comprehensive methodology would accurately reflect the physiological state of the baroreflex branches and their responses to orthostatic challenges.

## 2. METHODS

### 2.1. Study’s Subjects

We used two data sets from independent previous studies in the present study (Laude et al., 2004; Suarez-Roca et al., 2024). The first data set consisted of 55 patients scheduled for elective minimally invasive cardiothoracic surgery enrolled at Duke University Medical Center. These patients had high-quality blood pressure recordings and eurythmic electrocardiograms (ECG). The sample had a mean age of 63 years (SD = 9) and was 45% female. Of these patients, 27 (49%) underwent cardiac surgeries (minimally invasive valve repair or replacement with cardiopulmonary bypass). The remaining 28 patients (51%) underwent thoracic surgeries, including wedge resection, segmentectomy, or lobectomy performed via video-assisted thoracoscopic surgery (VATS).

Patients were excluded if they had preexisting persistent pain conditions (defined as pain lasting more than three months prior to enrollment), pain at the time of the interview with a PEG scale score > 1 or required preoperative opioid analgesics. Other exclusion criteria included a preexisting history of chronic atrial fibrillation or other arrhythmias with the use of Vaughan-Williams Class I or III antiarrhythmic medications, history of symptomatic cerebrovascular disease (e.g., prior stroke with residual deficits), alcoholism (defined as more than two drinks per day), psychiatric illness (any clinical diagnoses requiring therapy), drug abuse (any illicit drug use in the three months preceding surgery), severe pulmonary insufficiency (requiring home oxygen therapy), hepatic insufficiency (liver function tests ALT/AST > 1.5 times the upper limit of normal), and renal failure (serum creatinine > 2.0 mg/dl). This study was approved by the Institutional Review Board of Duke University Health System (IRB # Pro00083136), and written informed consent was obtained from all participants.

In addition, we used the EUROBAVAR dataset to evaluate the impact of positional changes on sympathetic baroreflex sensitivity (BRS). The EUROBAVAR dataset is an open-access collection of cardiovascular recordings from a *European multicenter study on baroreflex sensitivity and autonomic function in various diseases*, approved by the Paris-Necker Committee for the Protection of Human Subjects in Biomedical Research (available at http://www.eurobavar.altervista.org/). The EUROBAVAR dataset comprises continuous noninvasive blood pressure recordings in both supine and standing positions; it has been widely used to validate new methodologies and identify baroreflex patterns. The EUROBAVAR dataset consists of recordings from 21 subjects, including 17 women and 4 men, with a mean age of 38.4 years (SD = 15) (Laude et al., 2004). The cohort included 12 normotensive outpatients (one diabetic patient without cardiac neuropathy, two treated hypercholesterolemic subjects, and one 3-month pregnant woman), one untreated hypertensive, two treated hypertensive subjects, and four healthy volunteers. Notably, the EUROBAVAR dataset includes two baroreflex-impaired subjects (B005 and B010) due to diabetic autonomic neuropathy and a recent heart transplant procedure (Chao-Ecija et al., 2023) who were excluded from our present analysis. We employed the EUROBAVAR curve series format of the data, imported it into AcqKnowledge 5 software, and reconstructed the signal. This process allowed for detailed hemodynamic parameter analysis through contour analysis.

### 2.2. Noninvasive Collection and Processing of Cardiovascular Data

For the preoperative data set, the blood pressure and ECG signals were collected in a quiet, light-attenuated room with an ambient temperature of approximately 24°C. Before each evaluation, participants lay down in a semi-supine position (with the head of the bed elevated about 45 degrees) quietly for 10 minutes to allow their cardiovascular system to reach a steady state. Beat-by-beat arterial blood pressure was continuously recorded for 12-15 minutes using a double finger cuff device NIBP100D-HD (CNSystem, Graz, Austria), which enables non-disruptive arterial pressure recording in the finger based on the vasomotor unloading technique (Fortin et al., 2006). Simultaneously, ECG was recorded using a standard Lead II configuration. The biopotential signals were acquired and processed by the BIOPAC MP160/AcqKnowledge 5 system (BIOPAC Systems Inc., Goleta, CA, USA) with a sampling rate of 1 kHz.

For the EUROBAVAR data, the blood pressure and ECG signals were sampled at 500 Hz with a 16-bit resolution using the Finapres 2300 device (Ohmeda, Helsinki, Finland) and the Datex Cardiopac II monitor (Datex Engstrom, Helsinki, Finland) over 10-12 minutes in both supine and standing positions. The blood pressure and ECG biosignals from both datasets were passed through a Finite Impulse Response (FIR; a linear phase filter) digital filter with a 55-65 Hz band stop to remove 60 Hz electrical noise, ensuring no phase distortion between the original and filtered waveforms. The filtered signals were then subjected to artifact detection and cleaning.

Basic hemodynamic parameters, including systolic arterial pressure (SAP), diastolic arterial Pressure (DAP), mean arterial pressure (MAP), interbeat interval (IBI), heart rate, and the peak first derivative of the arterial pressure waveform (dP/dt_max_), were extracted from the pressure waveform and ECG recordings using AcqKnowledge software 5 with the ABP Classifier/Arterial Blood Pressure algorithm (BIOPAC Systems Inc., Goleta, CA, USA). Two hemodynamic variables were derived from these primary parameters: cardiac output (CO) and systemic vascular resistance (SVR). CO was noninvasively calculated by the CNAP® Monitor HD-500 using its built-in proprietary algorithm based on pulse contour analysis, which has an accuracy comparable to the gold-standard invasive thermodilution and PiCCO methods (Wagner et al., 2018). As a marker of afterload or arterial flow resistance, SVR (dyn.sec/cm^5^) = 80 x (MAP - CVP)/CO. For simplicity and consistency, we assumed a constant resting central venous pressure (CVP) of 0 mmHg, despite normal CVP typically ranging from 2 to 8 mmHg.

### 2.3. Assessment of Baroreflex Function: Components, Metrics, and Analytical Methods

We assessed five components of the baroreflex arc: the physiological stimulus (SAP) and four effector responses: sympathetically mediated peripheral vascular tone (estimated by SVR), myocardial contraction (estimated by arterial dP/dt_max_) (Morimont et al., 2012), the mixed sympathetic/parasympathetically regulated cardiac output, and parasympathetically mediated interbeat interval. For each effector response, we calculated three metrics: (1) baroreflex sensitivity (BRS), representing the gain of the reflex arc; (2) the baroreflex effectiveness index (BEI); and (3) the number of baroreflex sequences (SQ). These metrics were computed using a custom MATLAB algorithm based on the sequence method (Bertinieri et al., 1985; Parati et al., 2000b).

BRS was calculated by identifying baroreflex sequences with linear correlations greater than 0.9 between a stimulus sequence (at least three consecutive increases or decreases in SAP exceeding 0.5 units) and an effector response sequence (at least three consecutive changes in interbeat interval, dP/dt_max_, SVR exceeding 1 unit, or in CO greater than 0.1 unit). Positive correlations were considered for SAP with interbeat interval, and negative correlations for SAP with dP/dt_max_, SVR, and CO. BRS for each effector response was the arithmetic mean of individual BRS values from each baroreflex sequence.

Thus, we generated four BRS estimates: (1) the interbeat interval BRS (ibiBRS) was the slope of the change in interbeat interval per unit change in SAP (*ms/mmHg*); (2) the myocardial BRS (mBRS) was the slope of the change in dP/dt_max_ per unit change in SAP (*sec^-1^*); (3) the vasomotor BRS (vBRS) was the slope of the change in SVR per unit change in SAP (*dyn·s·cm^5^/mmHg*); and (4) the cardiac output BRS (coBRS) was the slope of the change in cardiac output per unit change in SAP (*ml/mmHg*) (Di Rienzo et al., 2001; Marchi et al., 2016). Similarly, we estimated BEI and SQ for each effector; BEI was the percentage of baroreflex sequences out of the total number of SAP ramps, and SQ was the number of baroreflex sequences over 10 minutes. We calculate these baroreflex metrics across 0, 1, 2, 3, and 6 beats delays relative to the blood pressure peak (SAP) to determine the influence of timing between the baroreflex stimulus and effector response.

### 2.4. Heart Rate Variability (HRV) Analysis: Time and Frequency Domain Measures

We analyzed ECG Lead II signals to extract standard HRV measures using AcqKnowledge5® based on guidelines from the European Society of Cardiology and the North American Society of Pacing and Electrophysiology (Berntson et al., 1997; Malik et al., 1996).

For HRV time domain analysis, we used the root mean square of the differences between successive R-R intervals (RMSSD; ms), which measures high-frequency heart rate variations and reflects parasympathetic heart regulation.

The frequency domain analysis involved (a) extracting RR intervals with AcqKnowledge 5’s QRS detector (BIOPAC Systems Inc., Goleta, CA, USA) using a modified Pan-Tompkins algorithm (Pan et al., 1985); (b) resampling RR intervals and applying cubic-spline interpolation for a continuous time-domain representation; and (c) generating power spectral density (PSD) from the interpolated tachogram using a fast Fourier transform-based Welch algorithm.

The analysis provided frequency domain measures in milliseconds squared (ms²) for three bands: high-frequency (HF-HRV, 0.15-0.4 Hz) reflecting parasympathetic activity, low-frequency (LF-HRV, 0.04-0.15 Hz) reflecting both sympathetic and parasympathetic activity (Topçu et al., 2018), and very-low-frequency (VLF-HRV, 0.0033-0.04 Hz) reflecting influences of several functions on heart rate, such as the renin-angiotensin system (Duprez et al., 1995; Taylor et al., 1998), vasomotor tone, peripheral chemoreceptor activity (Francis et al., 2000), and thermoregulation (Francis et al., 2000).

### 2.5. Data Analysis

Statistical analyses were conducted using SPSS for Windows, Version 29.0 (IBM Corp., Armonk, NY, USA). The significance level was set at α = 0.05 (two-sided). A repeated measures one-way ANOVA, followed by post hoc Bonferroni tests for multiple comparisons, was used to assess (1) the effect of beat delay on BRS, BEI, and SQ values and (2) the impact of positional changes and beat delay on these baroreflex metrics. A t-test was conducted to compare hemodynamic parameters between supine and standing positions.

Non-parametric Spearman’s correlation analysis determined the association between baroreflex parameters and hemodynamic variables. Values with Z-scores > 3 were considered outliers and excluded from the statistical analysis. The family-wise significance level was set at α = 0.05 (two-sided). Multiple testing corrections were performed using the Bonferroni method.

Statistical power was estimated using G*Power software (University of Duesseldorf, Germany) (Faul et al., 2007). With a sample size of 55, the study had sufficient power (Type II error < 0.2) to detect a correlation coefficient of 0.38 without multiple testing corrections, 0.45 when corrected for five multiple tests (P < 0.01), and 0.47 when corrected for ten multiple tests (P < 0.005). In the EUROBAVAR cohort (n = 21), a paired t-test would provide sufficient power (Type II error < 0.2) to detect an effect size of 0.65 between two means at P < 0.05 or an effect size of 0.82 at P < 0.01.

## 3. RESULTS

### 3.1. Validation of Cardiovascular Measurements through Correlation Analysis of Hemodynamic Variables and Baroreflex Components

We validated our cardiovascular measurements by identifying meaningful correlations between key hemodynamic variables and components of the baroreflex arc, averaged over 10 minutes. The analysis focused on the baroreflex stimulus (estimated by SAP) and the effector responses, which included peripheral vascular tone (estimated by SVR), myocardial contractility (estimated by dP/dt_max_ and cardiac output), and sinus atrial (SA) node activity (estimated as the interbeat interval).

After adjusting for multiple correlations, SAP levels (averaged over 10 minutes) exhibited significant positive correlations with other blood pressure parameters—diastolic arterial pressure (DAP), mean arterial pressure (MAP), and pulse pressure — as well as with dP/dt_max_ (Table 1). Additionally, SAP showed a positive correlation with interbeat interval and a negative correlation with heart rate (r = 0.406, p = 0.002). These correlations align with established physiological principles (Hall et al., 2011). In normotensive subjects, increased SAP raises DAP, MAP, and PP, triggering a compensatory decrease in heart rate via the baroreceptor reflex to stabilize blood pressure and reducing high-frequency heart rate variability power (HRV-HF), reflecting decreased parasympathetic activity.

**Table 1.** Correlations between Components of the Baroreflex Arc (Stimulus and Effector Responses) and Selected Hemodynamic Variables.

| Hemodynamics parameters | *Stimulus parameter* |  | *Sympathetically modulated response parameter* |  |  |  |  |  | *Parasympathetically modulated response parameter* |  |
| --- | --- | --- | --- | --- | --- | --- | --- | --- | --- | --- |
|  | Systolic | *p-value* | SVR | *p-value* | dP/dt<sub>max</sub> | *p-value* | CO | *p-value* | IBI | *p-value* |
| **Blood pressures** |  |  |  |  |  |  |  |  |  |  |
| SAP |  |  | 0.281 | 0.037 | **0.634\*** | <0.001 | 0.223 | 0.102 | **0.406\*** | 0.002 |
| DAP | **0.476\*** | <0.001 | 0.355 | 0.008 | -0.207 | 0.130 | 0.180 | 0.189 | 0.183 | 0.186 |
| MAP | **0.762\*** | <0.001 | 0.369 | 0.006 | 0.123 | 0.373 | 0.234 | 0.086 | 0.333 | 0.014 |
| PP | **0.636\*** | <0.001 | -0.038 | 0.781 | **0.900\*** | <0.001 | 0.117 | 0.395 | 0.302 | 0.026 |
| LVET | 0.311 | 0.021 | 0.085 | 0.537 | 0.091 | 0.508 | 0.087 | 0.526 | **0.405\*** | 0.002 |
| SAP-RMSSD | -0.110 | 0.423 | 0.118 | 0.389 | 0.034 | 0.806 | -0.371 | 0.005 | **-0.413\*** | 0.002 |
| **Inotropic** |  |  |  |  |  |  |  |  |  |  |
| dP/dt<sub>max</sub> | **0.634\*** | <0.001 | 0.006 | 0.967 | 1.000 |  | 0.109 | 0.430 | 0.213 | 0.121 |
| SV | 0.307 | 0.023 | **-0.583\*** | 0.001 | 0.169 | 0.218 | **0.861\*** | <0.001 | **0.722\*** | <0.001 |
| CO | 0.223 | 0.102 | **-0.761\*** | 0.001 | 0.109 | 0.430 |  |  | 0.339 | 0.012 |
| SVR | 0.281 | 0.037 |  |  | 0.006 | 0.967 | **-0.761\*** | <0.001 | -0.083 | 0.552 |
| **Chronotropic & Dromotropic** |  |  |  |  |  |  |  |  |  |  |
| IBI | **0.412\*** | 0.002 | -0.076 | 0.585 | 0.220 | 0.110 | 0.339 | 0.012 |  |  |
| QRS | 0.087 | 0.532 | -0.089 | 0.524 | **0.404\*** | 0.002 | -0.005 | 0.971 | -0.057 | 0.682 |
| PRI | 0.028 | 0.839 | 0.266 | 0.052 | 0.041 | 0.766 | -0.212 | 0.124 | 0.045 | 0.749 |
| **Heart rate variability** |  |  |  |  |  |  |  |  |  |  |
| RMSSD | -0.140 | 0.309 | 0.226 | 0.097 | -0.086 | 0.533 | -0.393 | 0.003 | 0.093 | 0.504 |
| Total Power | -0.049 | 0.721 | 0.143 | 0.298 | 0.064 | 0.641 | -0.295 | 0.029 | -0.054 | 0.696 |
| VLF | 0.037 | 0.790 | -0.231 | 0.090 | -0.064 | 0.643 | 0.201 | 0.141 | -0.199 | 0.149 |
| LF | 0.037 | 0.790 | -0.231 | 0.090 | -0.064 | 0.643 | 0.201 | 0.141 | -0.199 | 0.149 |
| HF | -0.360 | 0.007 | 0.058 | 0.673 | -0.042 | 0.762 | -0.285 | 0.035 | -0.169 | 0.222 |
| LF/HF ratio | 0.314 | 0.020 | -0.075 | 0.588 | -0.002 | 0.991 | 0.268 | 0.048 | 0.104 | 0.454 |

**Note:** The values of the hemodynamic variables used in the correlations are averaged over a 10-minute recording from preoperative patients (N = 55). Abbreviations: SAP = Systolic Arterial Pressure (mmHg); DAP = Diastolic Arterial Pressure (mmHg); MAP = Mean Arterial Pressure (mmHg); PP = Pulse Pressure (mmHg); LVET = Left ventricle ejection time; SAP-RMSSD = the Root Mean Square of Successive Differences between systolic arterial pressure (mmHg); dP/dt<sub>max</sub> = maximal rate of pulse pressure changes over time (mmHg/sec); CO = Cardiac Output (L/min); SV = Stroke Volume (ml), SVR (dynes-sec/cm<sup>5</sup>); IBI = Interbeat Interval (ms); QRS = ECG QRS complex duration (ms), PRI = ECG P-R Interval (ms); RMSSD = root mean square of successive differences of ECG RR intervals; Total Power = HRV-Total Spectral Power (Ln ms<sup>2</sup>); VLF = HRV very low-frequency power (Ln n.u.); LF = HRV low-frequency power (Ln n.u.); HF = HRV high-frequency power (Ln n.u.); LF/HF ratio = HRV-LF(n.u.) / HRV-HF(n.u.) (Ln). **In bold text:** \* Significant correlations between the stimulus or response and the 18 hemodynamic parameters; significance level adjusted for multiple comparisons with $\alpha = 0.05/18 = 0.0027$ (two-tailed).

Regarding baroreflex responses, SVR was negatively correlated with cardiac output and stroke volume **(Table 1)**, indicating that increased vascular resistance reduces cardiac efficiency. The dP/dt_max_ was positively correlated with SAP, PP, and QRS duration, linking stronger and longer myocardial contractions with higher blood pressure. As expected, cardiac output was positively correlated with stroke volume and negatively correlated with SVR (Reddy et al., 2016). Finally, interbeat interval showed positive correlations with SAP, left ventricular ejection time (LVET), and stroke volume while negatively correlating with SAP variability (estimated as SAP-RMSSD), which is consistent with known baroreflex-mediated associations (Parati et al., 2000a; Ursino, 1999). These correlations are physiologically consistent and meaningful, validating the accuracy of our hemodynamic and ECG estimations.

### 3.2. Estimation of Baroreflex Parameters for Discrete Effectors Across Beat Delays

**Figure 1** presents the analysis of how four different baroreflex effectors respond to changes in SAP across various beat delays (measured in heartbeats). A repeated measure one-way ANOVA applied to each baroreflex metric—BRS, BEI, and SQ—for each baroreflex effector revealed distinctive response patterns. BRS remained consistent across beat delays, showing no significant changes for any of the four effectors. In contrast, BEI and SQ metrics exhibited significant variations depending on the beat delay, with these changes being specific to each effector (i.e., peripheral vascular tone, myocardial contractility, cardiac output, or heart rate).

**Figure 1.**
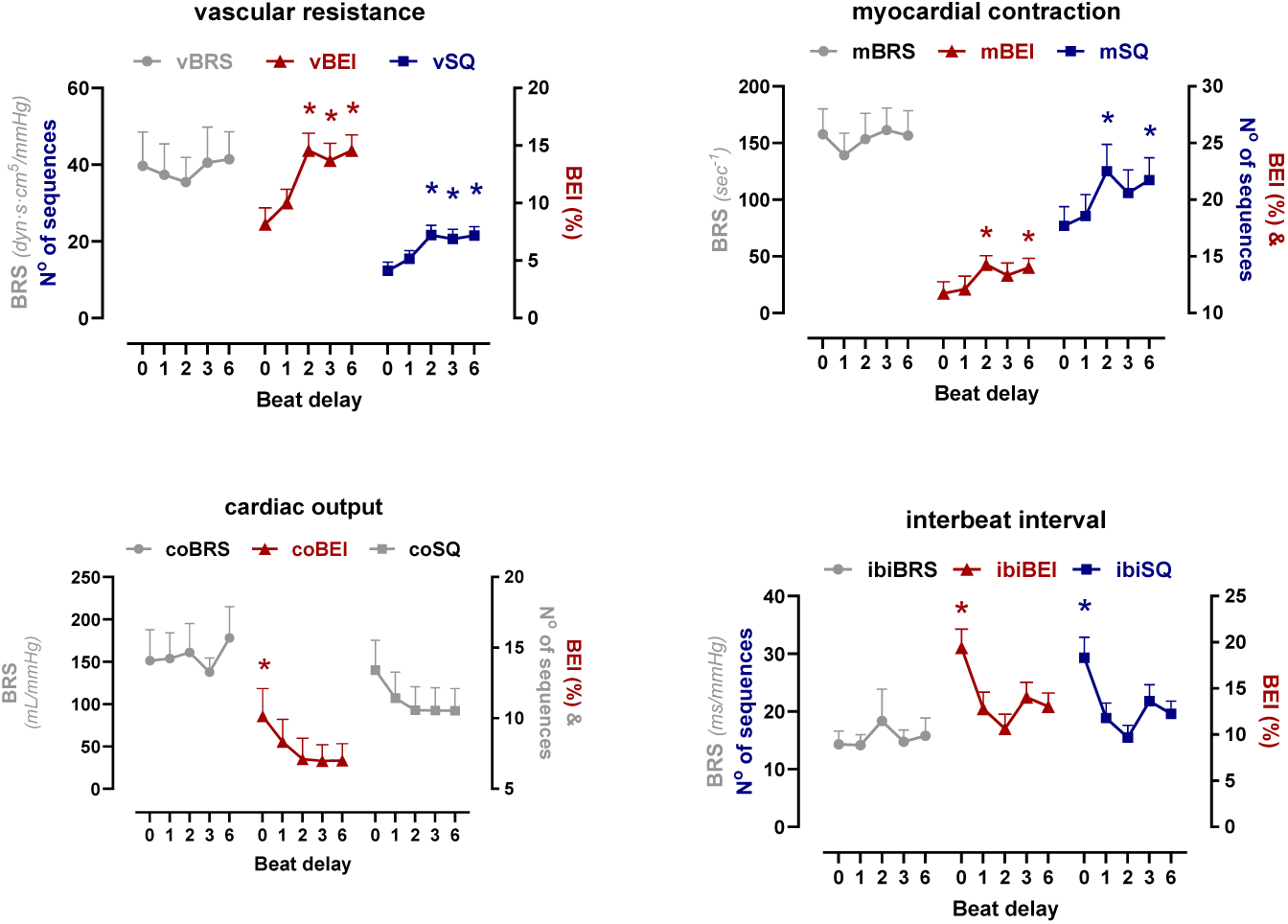
Influence of systolic arterial pressure (SAP) on four baroreflex effectors: peripheral vascular tone (SVR), myocardial contraction (dP/dtmax), cardiac output (CO), and heart rate (HR) across different beat delays (0, 1, 2, 3, and 6). Each panel displays three parameters: baroreflex sensitivity (BRS, circles), baroreflex effectiveness index (BEI, triangles), and the number of baroreflex sequences over 10 minutes (SQ, squares). Data symbols represent the mean ± 95% confidence interval. Colored lines contain statistically significant values, with asterisks denoting significant differences compared to the lowest data point in the line (repeated measures one-way ANOVA, followed by Bonferroni-corrected post hoc tests for multiple comparisons; p < 0.05, two-tailed test).

For vascular resistance, both BEI (vBEI) and SQ (vSQ) values significantly increased at beat delays of 2, 3, and 6 compared to the 0-beat delay, which was the lowest point (F _(4, 268)_ = 17.721, p < 0.001; F _(4, 267)_ = 12.805, p < 0.001; respectively). This pattern suggests that the sympathetic baroreflex-mediated vascular response is not immediate but shows a sustained increase following the initial change in SAP. Significant differences in BEI and SQ were also observed for myocardial contraction (mBEI and mSQ) at beat delays of 2 and 6 (F _(4, 270)_ = 5.226, p < 0.001; F _(4, 270)_ = 4.208, p = 0.003; respectively); this implies that myocardial responses to sympathetic baroreflex activation may be optimally elicited or more detectable after a slight delay from the initial SAP change. For cardiac output, a statistically significant peak in BEI (coBEI) was observed at a 0-beat delay (F _(4, 269)_ = 3.331, p = 0.008), though there was no significant change in SQ (coSQ) (F _(4, 268)_ = 1.963, p = 0.100). Finally, the BEI and SQ values for interbeat interval responses (hrBEI and hrSQ) were highest at the 0-beat delay (F _(4, 269)_ = 14.781, p < 0.001; F _(4, 268)_ = 14.425, p < 0.001), indicating that the heart rate response to baroreflex activation is most immediate.

### 3.3. BRS Associations with Average Levels of the Hemodynamic and HRV Parameters

BRS for systemic vascular resistance (vBRS) showed significant correlations with the average values of baroreflex hemodynamic stimuli and responses across all delays, particularly after 2 or 3 beats from the SAP peak **(Figure 2)**. Specifically, vBRS negatively correlated with the ongoing levels of SAP (averaged over 10 minutes) at a 3-beat delay, even after adjusting for multiple comparisons (p < 0.01). Additionally, vBRS demonstrated strong correlations with vascular and inotropic parameters across various delays. It positively correlated with average SVR values at the 0-beat delay and after 2, 3, and 6 beats from the SAP peak. Conversely, vBRS negatively correlated with cardiac output across all delays and with dP/dt_max_ at a one-beat delay (p < 0.01). As for chronotropic parameters, vBRS negatively correlated with average interbeat interval values at a 6-beat delay (p < 0.01). Regarding cardiac autonomic modulation, vBRS positively correlated with RMSSD-HRV at 0-, 2-, and 3-beat delays (p < 0.01).

**Figure 2.**
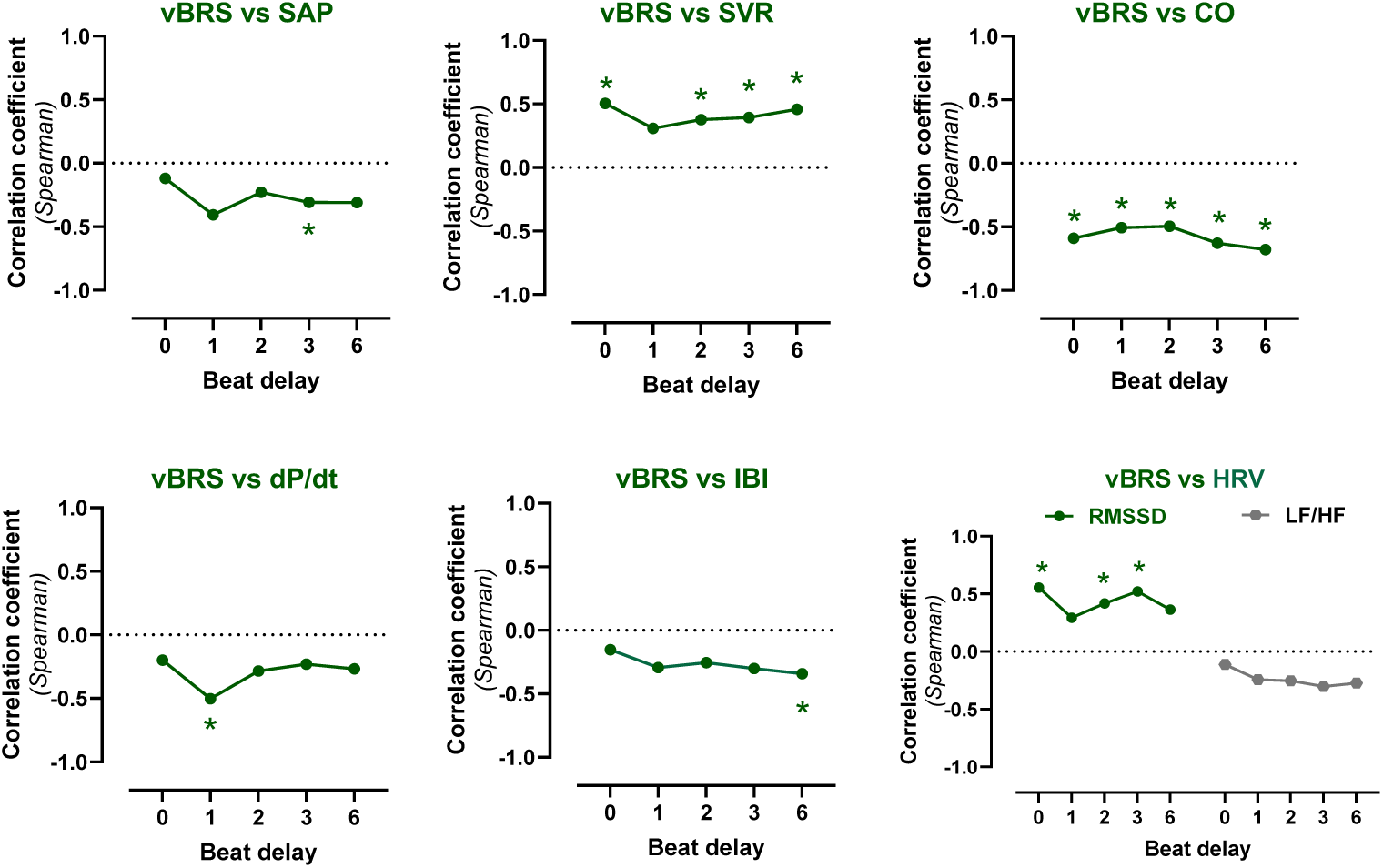
Correlations of vascular BRS (vBRS) with hemodynamic and HRV parameters across beat delays relative to the SAP peak. The hemodynamic parameters analyzed include systolic arterial pressure (SAP), systemic vascular resistance (SVR), cardiac output (CO), maximal rate of pulse pressure changes over time (dP/dt), interbeat interval (IBI), root mean square of successive differences in heart rate (RMSSD), and the low-frequency to high-frequency HRV ratio (LF/HF). Dots represent Spearman’s correlation coefficients. *Significant correlations are indicated at five beat delays; the significance level was adjusted for multiple comparisons with α = 0.01 (two-tailed test). For correlations involving five beat delays and two HRV parameters (RMSSD and LF/HF), the significance level was further adjusted to α = 0.005 (two-tailed test).

**Figure 3** illustrates that BRS for modulating myocardial contraction (mBRS) positively correlated with average dP/dt_max_ at a one-beat delay (p < 0.01). mBRS did not correlate with any other hemodynamic or HRV parameters.

**Figure 3.**
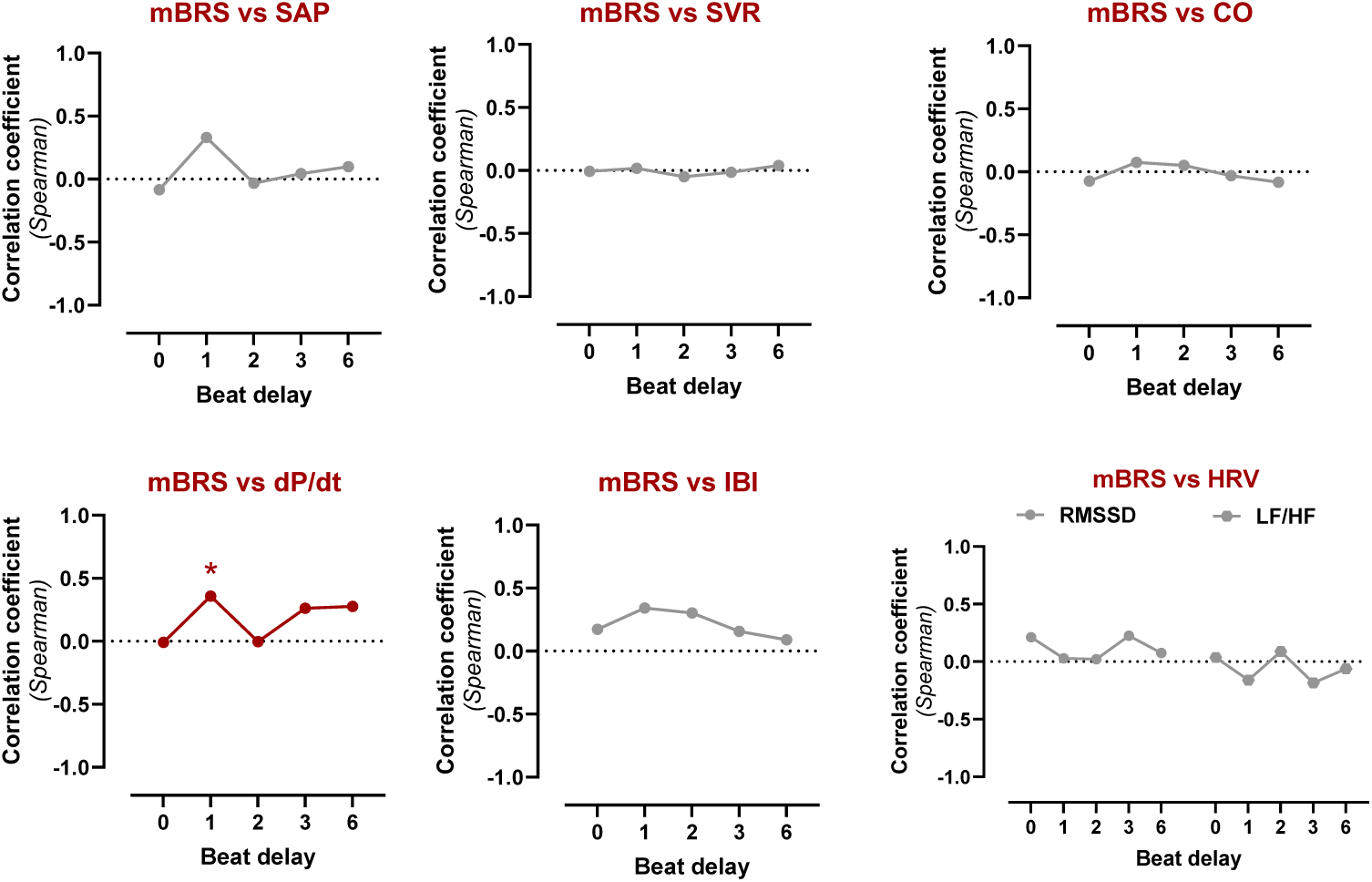
Correlations of myocardial BRS (mBRS) with hemodynamic and HRV parameters across beat delays relative to the SAP peak. The hemodynamic parameters analyzed include systolic arterial pressure (SAP), systemic vascular resistance (SVR), cardiac output (CO), maximal rate of pulse pressure changes over time (dP/dt), interbeat interval (IBI), root mean square of successive differences in heart rate (RMSSD), and the low-frequency to high-frequency HRV ratio (LF/HF). Dots represent Spearman’s correlation coefficients. *Significant correlations are indicated at five beat delays; the significance level was adjusted for multiple comparisons with α = 0.01 (two-tailed test).

BRS for modulating cardiac output (coBRS) was correlated with the average levels of cardiac output at 0- and 3-beat delays **(Figure 4)**. Additionally, coBRS negatively correlated with average SVR values at 0-, 1-, 2-, and 3-beat delays (p < 0.01) and positively with cardiac output at 0- and 3-beat delays (p < 0.01). However, coBRS did not correlate with other hemodynamic and HRV parameters.

**Figure 4.**
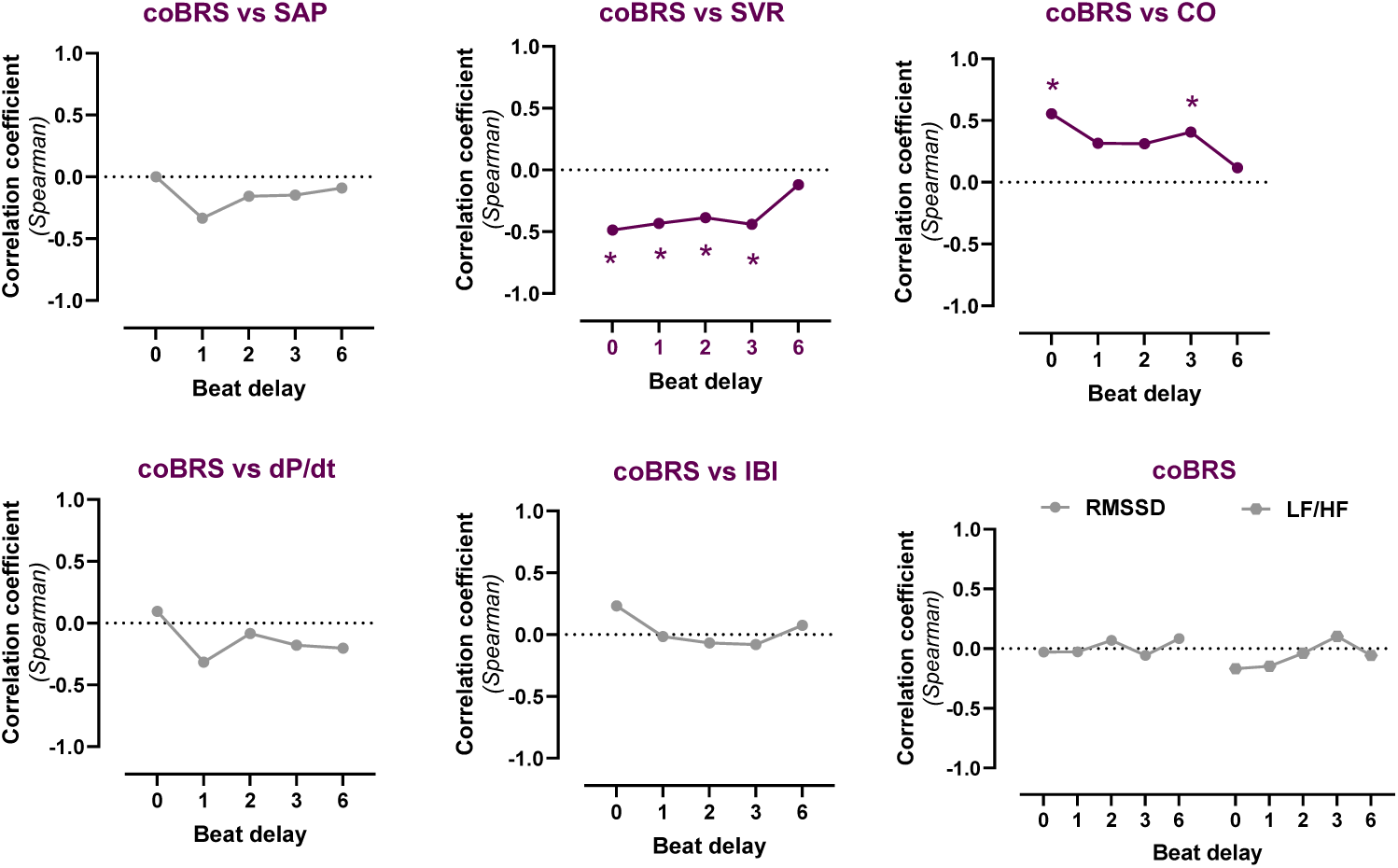
Correlations of BRS for cardiac output (coBRS) with hemodynamic and HRV parameters across beat delays relative to the SAP peak. The hemodynamic parameters analyzed include systolic arterial pressure (SAP), systemic vascular resistance (SVR), cardiac output (CO), maximal rate of pulse pressure changes over time (dP/dt), interbeat interval (IBI), root mean square of successive differences in heart rate (RMSSD), and the low-frequency to high-frequency HRV ratio (LF/HF). Dots represent Spearman’s correlation coefficients. *Significant correlations are indicated at five beat delays; the significance level was adjusted for multiple comparisons with α = 0.01 (two-tailed test).

BRS for modulating the interbeat interval (ibiBRS) assessed the strength of inhibitory baroreflex modulation on heart rate, as reflected in the prolongation of the IBI. ibiBRS positively correlated with average RMSSD-HRV at the 0-beat delay (p < 0.01) **(Figure 5)**. It also negatively correlated with average cardiac output values at a 3-beat delay (p < 0.01) but showed no correlation with SVR. ibiBRS was not associated with other hemodynamic or HRV parameters.

**Figure 5.**
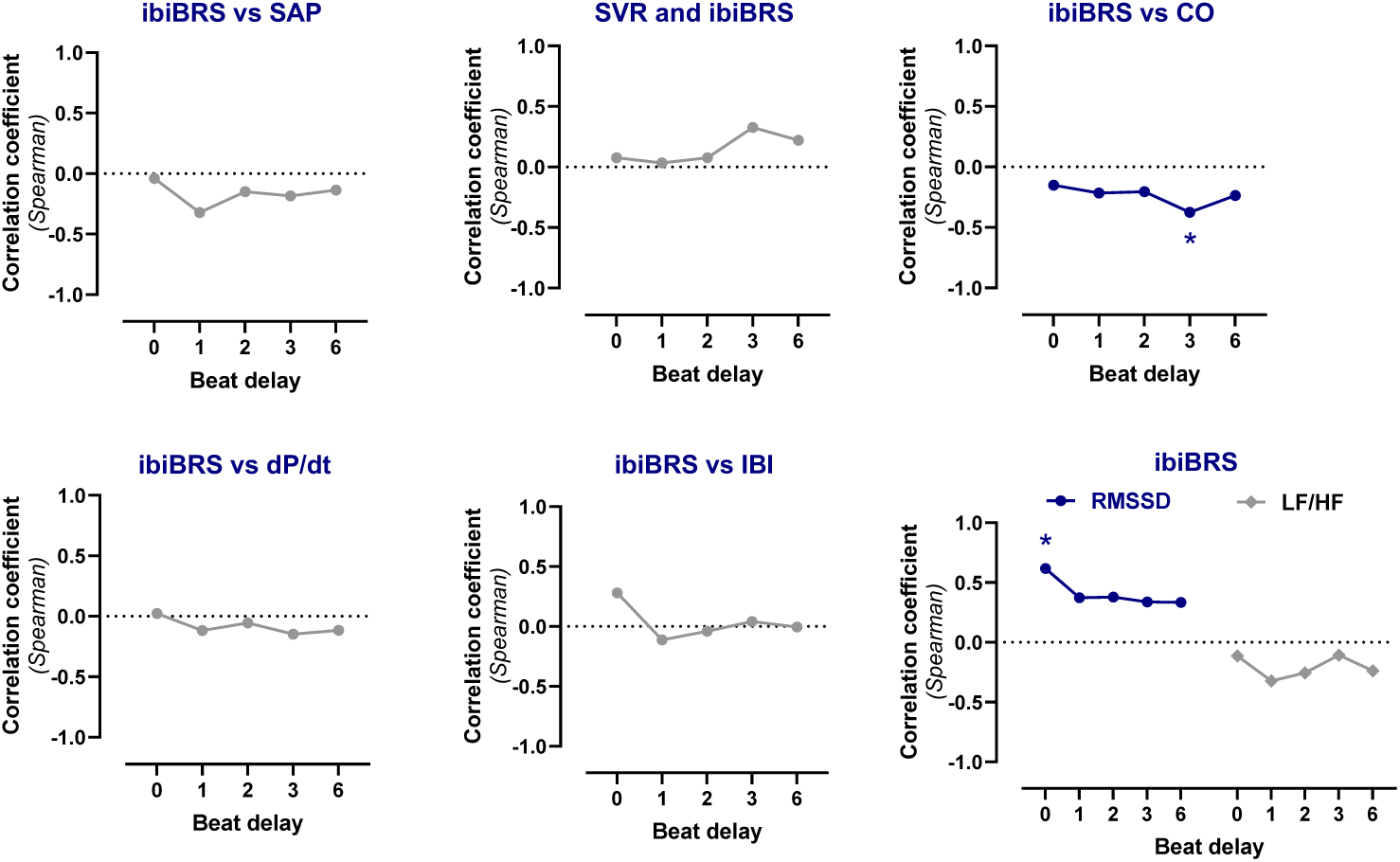
Correlations of BRS for interbeat interval (ibiBRS) with hemodynamic and HRV parameters across beat delays relative to the SAP peak. The hemodynamic parameters analyzed include systolic arterial pressure (SAP), systemic vascular resistance (SVR), cardiac output (CO), maximal rate of pulse pressure changes over time (dP/dt), interbeat interval (IBI), root mean square of successive differences in heart rate (RMSSD), and the low-frequency to high-frequency HRV ratio (LF/HF). Dots represent Spearman’s correlation coefficients. *Significant correlations are indicated at five beat delays; the significance level was adjusted for multiple comparisons with α = 0.01 (two-tailed test). For correlations involving five beat delays and two HRV parameters (RMSSD and LF/HF), the significance level was further adjusted to α = 0.005 (two-tailed test).

### 3.4. Cardiovascular Adaptations to Postural Changes

**Table 2** shows that SAP, MAP, and PP did not significantly change in response to postural changes. Similarly, dP/dtmax, cardiac output, and SVR remained stable. In contrast, heart rate and DAP, averaged over 10 minutes, increased from 72 to 83 bpm during standing, while left ventricular ejection time (LVET) decreased. Regarding autonomic modulation, RMSSD-HRV decreased, and the LF/HF-HRV ratio increased during standing compared to the supine position.

**Table 2:** Cardiovascular Adaptations to Postural Changes from Supine to Standing Positions.

| Cardiovascular parameters | Supine |  | Standing |  | p-value |
| --- | --- | --- | --- | --- | --- |
|  | Mean | SD | Mean | SD |  |
| Systolic arterial pressure (*mmHg*) | 120.5 | 18.7 | 121.2 | 21.4 | 0.7990 |
| **Diastolic arterial Pressure (*mmHg*)** | 61.4 | 11.7 | 67.9 | 11.6 | **<0.0001\*** |
| Mean arterial pressure (*mmHg*) | 81.1 | 13.7 | 85.6 | 14.2 | 0.0069 |
| Pulse pressure (*mmHg*) | 58.9 | 9.7 | 53.2 | 13.5 | 0.0100 |
| dP/dt<sub>max</sub> (*mmHg/sec*) | 1494 | 283 | 1592 | 392 | 0.1958 |
| **Left ventricle ejection time (*ms*)** | 206 | 65 | 184 | 48 | **0.0016\*** |
| Cardiac output (*L/min*) | 5.09 | 2.09 | 6.47 | 4.18 | 0.2378 |
| Systemic vascular resistance (*dynes·sec/cm<sup>5</sup>*) | 1718 | 1501 | 1289 | 608 | 0.2457 |
| **Heart rate (*bpm*)** | 72.1 | 13.6 | 83.1 | 15.1 | **<0.0001\*** |
| **RMSSD-HRV (*ms*)** | 37.6 | 29.2 | 23.1 | 16.6 | **0.0009\*** |
| **LF/HF-HRV ratio (*n.u.*)** | 1.35 | 1.07 | 1.92 | 1.76 | **0.0009\*** |

**Note:** Cardiovascular parameters are presented as beat-to-beat values averaged over a 10-minute recording. \*Significant p-values after adjusting for ten hemodynamic parameters, with a significance level of $\alpha = 0.005$ (paired t-test, two-tailed; N = 21). maximal rate of pulse pressure changes over time (dP/dt<sub>max</sub>), RMSSD-HRV = root mean square of successive differences of ECG RR intervals. LFnu/HFnu HRV ratio = Low-frequency to high-frequency HRV ratio in normalized units. RMSSD-HRV and the LF/HF-HRV ratio were logarithmically transformed (Ln) for statistical analysis.

### 3.5. Impact of Postural Changes on Baroreflex Parameters Across Beat Delays

We evaluated the impact of postural changes—from supine to standing—on baroreflex effectiveness (BEI) and the number of baroreflex sequences (SQ) across beat delays in the EUROBAVAR group (**Figure 6**). Postural changes notably affected the baroreflex regulation of SVR. During standing, vBEI and vSQ reached their highest values at beat delays of 3 and 6 (F _(4, 100)_ = 8.860, p < 0.001; F _(4, 100)_ = 9.213, p < 0.001; respectively), which is considerably later than the peak values observed in the supine position at 1 and 2 beat delays (F _(4, 100)_ = 11.927, p < 0.001; F _(4, 100)_ = 8.633, p < 0.001; respectively).

**Figure 6.**
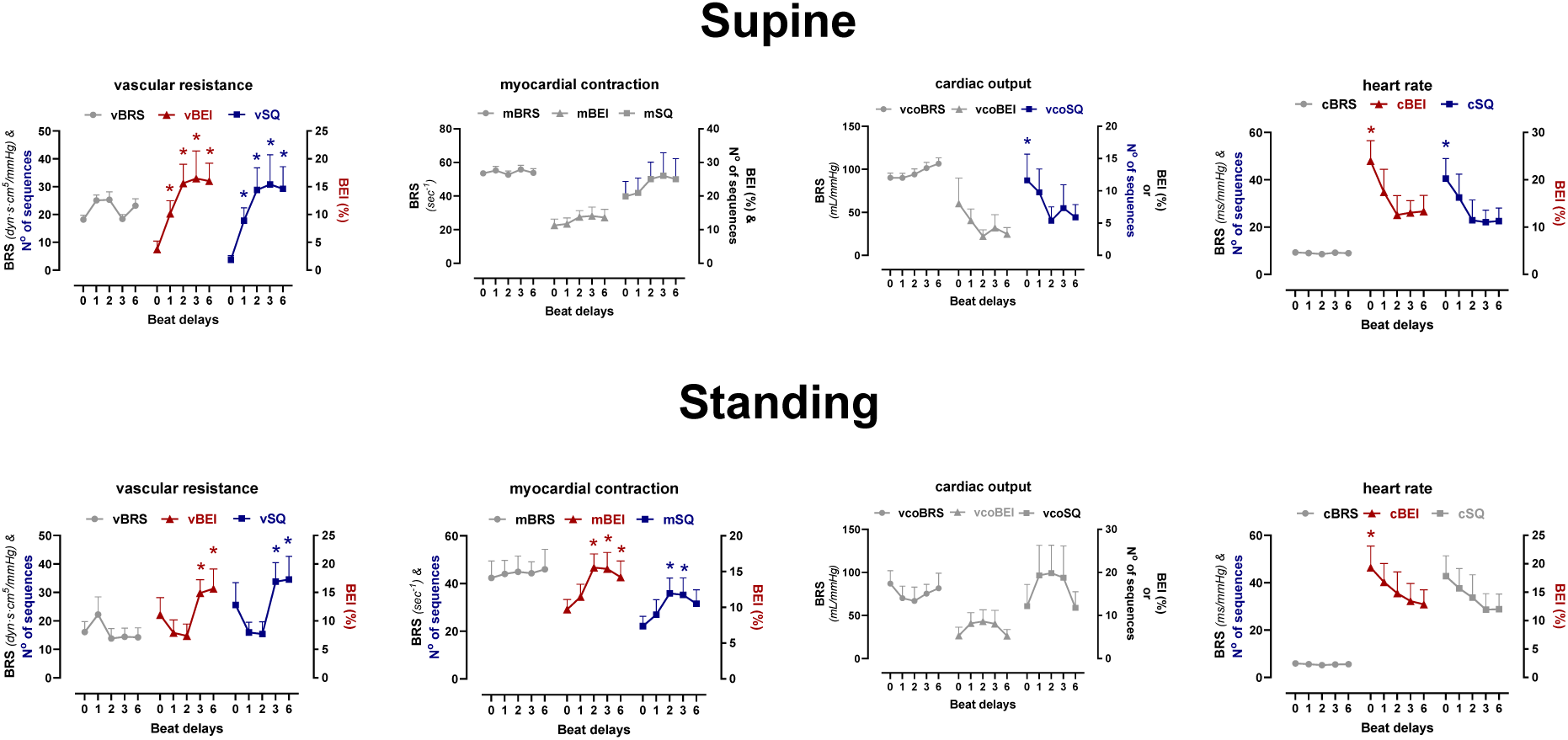
Impact of Postural Changes on Peak BRS, BEI, and SQ Values. This figure illustrates the effects of postural changes— from supine to standing—on various types of baroreflex sensitivity (BRS): vascular (vBRS), myocardial (mBRS), cardiac output (coBRS), and heart rate (ibiBRS) across beat delays of 0, 1, 2, 3, and 6. Each panel shows BRS values (mean ± 95% confidence interval). Colored lines contain statistically significant values, with asterisks denoting significant differences compared to the lowest data point in the line (repeated measures one-way ANOVA, followed by Bonferroni-corrected post hoc tests for multiple comparisons; p < 0.05, two-tailed test).

Similarly, the effectiveness and frequency of baroreflex regulation of myocardial contraction, as estimated by mBEI and mSQ, were significantly higher at late beat delays (2, 3, and 6) during standing (F _(4, 100)_ = 7.720, p < 0.001; F _(4, 100)_ = 7.720, p < 0.001; respectively). In contrast, in the supine position, mBEI and mSQ did not show significant differences across beat delays (F _(4, 100)_ = 1.626, p = 0.173; F _(4, 100)_ = 1.119, p = 0.352; respectively), which may be attributed to the smaller sample size in this group.

For cardiac output regulation, the highest values for coBEI and coSQ were observed at 0 beat delay in the supine position (F _(4, 100)_ 3.653, p = 0.008; F _(4, 100)_ = 2.972, p = 0.023, respectively), but these values did not reach significance across any beat delay during standing (F _(4, 100)_ 2.069, p = 0.090; F _(4, 100)_ = 1.930, p = 0.111, respectively). This difference may be due to the smaller sample size in the standing group.

Regarding heart rate regulation, the highest values for hrBEI and hrSQ were observed at 0 beat delay and were not significantly affected by postural changes (standing: hrBEI - F _(4, 100)_ = 2.698, p = 0.035; hrSQ - F _(4, 100)_ = 2.440, p = 0.052; supine: hrBEI - F _(4, 100)_ = 6.548, p = 0.000; hrSQ - F _(4, 100)_ = 4.897, p = 0.001).

A repeated measures two-way ANOVA revealed that BRS for modulating myocardial contraction (mBRS) and interbeat interval (ibiBRS) were reduced during standing compared to the supine position (**Figure 7**). Specifically, mBRS was significantly lower during standing at 0 beat delay (F _(1, 36)_ = 4.512, p= 0.0406), although the beat delay x position interaction was not significant (F _(4, 143)_ = 0.7234, p = 0.5773), indicating consistent postural effects across beat delays. Similarly, ibiBRS was consistently lower in the standing position across all delays—i.e., 0, 1, 2, 3, and 6 beats—with p-values of 0.0066, 0.0029, 0.0006, 0.0005, and 0.0048, respectively (F _(1, 35)_ = 13.46, p=0.0008). The position x beat delay interaction was not significant (F _(4, 140)_ = 0.1509, p=0.9623).

**Figure 7.**
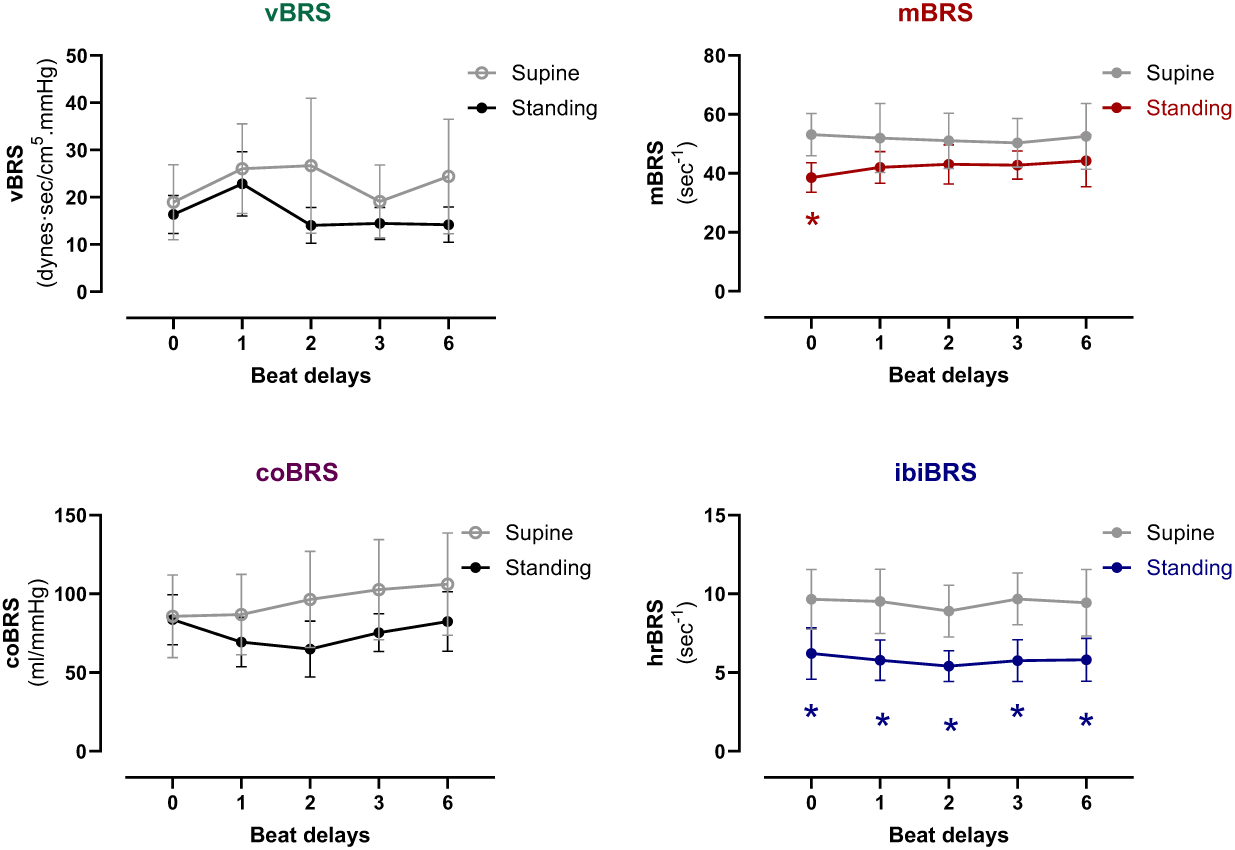
Impact of Postural Changes on Different BRS Types. This figure illustrates the effects of postural changes—from lying down to standing—on various types of baroreflex sensitivity (BRS): vascular (vBRS), myocardial (mBRS), cardiac output (coBRS), and interbeat interval (ibiBRS) across beat delays of 0, 1, 2, 3, and 6. Each panel shows BRS values (mean ± 95% confidence interval) in both lying down (gray lines) and standing (black, red, and blue lines) positions. Red and blue lines in the standing position indicate a statistically significant main effect of position, highlighting differences compared to the corresponding beat delay value in the lying down position (repeated measures two-way ANOVA, followed by Bonferroni-corrected post hoc tests for multiple comparisons; p < 0.05, two-tailed test).

The analysis did not show a significant main effect of position or beat delay x position interaction on vBRS or coBRS.

## 4. DISCUSSION

This study developed a simple, noninvasive method for simultaneously assessing the temporal dynamics of multiple baroreflex branches by estimating beat-to-beat changes in SVR, dP/dt_max_, cardiac output, and interbeat interval in response to SAP oscillations using pulse wave contour analysis.

The sequence method enabled the calculation of BRS, BEI, and SQ at various beat delays. Unlike comparable methods, our approach consistently detected numerous reflex sequences for each baroreflex branch.

We validated this method by demonstrating significant correlations between beat-to-beat BRS and SAP, SVR, dP/dt_max_, cardiac output, and interbeat interval averaged over 10 minutes. These correlations depended on beat delay and were consistent with known physiological interrelationships and autonomic response time frames.

### 4.1. Autonomic Nature of Baroreflex-Mediated Responses Identified by Beat Delays

BRS values remained stable across beat delays, suggesting consistent baroreflex gain regardless of the autonomic mechanism. However, BEI and SQ values varied, with the highest observed at 0-beat delay for ibiBRS and coBRS and between 2 to 6-beat delays for vBRS and mBRS. These patterns reflect differences in response latencies between sympathetic and parasympathetic pathways (Reyes Del Paso et al., 2017). In humans, the inhibition of the peroneal muscle sympathetic nerve occurs 1.16 – 1.49 seconds after the QRS complex of the ECG (Sundlof et al., 1978), and sympathetic bursts arise 1.20 – 1.40 seconds after the DAP trough (Kienbaum et al., 2001). In dogs, perfusion pressure in resistance arteries rises within 0.2 – 2.0 seconds after sympathetic nerve stimulation and falls within 0.5 – 5 seconds after stimulation ends (Rosenbaum et al., 1968). These findings suggest that sympathetically-mediated baroreflex responses likely occur between 2 and 8 beats after the SAP peak in subjects with an average heart rate.

### 4.2. Vascular BRS (vBRS) and Its Associations with Hemodynamic and HRV Parameters

vBRS reflects the strength of baroreflex inhibition on sympathetically-regulated vascular resistance. The maximal vascular baroreflex effectiveness (vBEI) and frequency (vSQ) were observed at 2 to 6-beat delays, indicating that the vascular baroreflex predominantly operates within these latencies, consistent with the time frame known for sympathetically mediated vascular baroreflexes (Kienbaum et al., 2001; Rosenbaum et al., 1968). In line with this, subjects with elevated vBRS exhibited increased average SVR levels and, correspondingly, a smaller average cardiac output across the same 2 to 6-beat delays. Interestingly, higher vBRS was related to lower SAP levels at a 3-beat delay. Notably, this reciprocal relationship between SAP and vBRS was observed exclusively with vBRS but not with other baroreflex branches. These findings suggest that vBRS might serve as a compensatory mechanism, inhibiting sympathetic vascular tone to maintain a normal SAP under elevated SVR conditions.

Higher vBRS negatively correlated with average dP/dt_max_ at a one-beat delay. This negative correlation could result from the increased SVR present at this latency, leading to an elevated afterload and slower myocardial contraction (Monge Garcia et al., 2018), which may account for the observed reduction in cardiac output. However, this association might be physiologically irrelevant, as baroreflex effectiveness and frequency were very low at this latency.

Furthermore, higher vBRS was also related to a lower average interbeat interval (i.e., higher heart rate) at the 6-beat delay, which may reflect a secondary response to a diminished cardiac output. Additionally, higher vBRS was linked to an increased RMSSD-HRV at a 0-beat delay and after 2- and 3-beats, suggesting the involvement of early parasympathetic and late sympathetic mechanisms.

### 4.3. Myocardial BRS and Its Associations with Hemodynamic and HRV Parameters

mBRS represents the inhibitory baroreflex modulation of sympathetically regulated myocardial contractility. Similar to the vascular baroreflex, the maximal myocardial baroreflex effectiveness (vBEI) and frequency (vSQ) were observed at 2 to 6-beat delays, indicating the involvement of sympathetic mechanisms (Kienbaum et al., 2001; Rosenbaum et al., 1968). Subjects with higher mBRS exhibited faster ventricular contraction—average dP/dt_max_— at a one-beat delay. This finding suggests that higher mBRS likely reflects a homeostatic response aimed at reducing elevated myocardial contraction by inhibiting the sympathetic drive. Notably, this sympathetic response is faster than that observed in the vascular baroreflex (2-3 beat delay), likely due to the shorter neural pathways involved (Charkoudian et al., 2009). However, the myocardial baroreflex on dP/dt_max_ may be relatively weak, as its effectiveness and frequency were very low at this latency.

Furthermore, mBRS was specific for assessing baroreflex modulation of heart contractility, showing no significant associations with other baroreflex responses or HRV parameters. An advantage of mBRS, based on beat-to-beat dP/dt_max_ response to SAP change, is that it avoids the confounding influence of heart rate and the limited baroreflex sequences observed with stroke volume or systolic pre-ejection periods (Reyes Del Paso et al., 2017).

### 4.4. BRS for Cardiac Output and Its Associations with Hemodynamic and HRV Parameters

coBRS estimates the strength of baroreflex inhibition on cardiac output under the influence of both sympathetic and parasympathetic drives, which regulate stroke volume and heart rate, respectively (Hall et al., 2011). Subjects with higher coBRS exhibited elevated average cardiac output and a diminished average SVR at 0-beat and 3-beats after the SAP peak, consistent with dual autonomic regulation. Notably, the maximal baroreflex effectiveness on cardiac output (coBEI) was observed at a 0-beat delay, suggesting that the fast parasympathetic control is predominant over the slow sympathetic drive. The sympathetic component becomes evident during low-intensity exercise, where the carotid sinus baroreflex maintains blood pressure despite a reduced BRS for heart rate due to parasympathetic blockade (Fisher et al., 2006).

The associations between coBRS, SVR, and cardiac output mirrored those seen with vBRS, indicating an antagonistic dynamic between these two baroreflex branches. An increased coBRS might counteract elevated cardiac output resulting from reduced afterload secondary to diminished SVR (Reddy et al., 2016).

### 4.5. BRS for Heart Rate and Its Associations with Hemodynamic and HRV Parameters

Although ibiBRS, which measures the inhibitory baroreflex modulation of heart rate, was not associated with average interbeat intervals, it showed a positive correlation with RMSSD-HRV at a 0-beat delay, a time frame indicative of fast parasympathetic modulation. Consistently, improvements in cardiovagal BRS are linked to increased HRV in humans (Shaltout et al., 2018; Suarez-Roca et al., 2024). Conversely, higher ibiBRS was associated with lower average cardiac output at a 3-beat delay, suggesting delayed baroreflex inhibition of sympathetic input to the heart (Karemaker, 2022).

### 4.6. Comparative Analysis of Sympathetic Baroreflex Mechanisms

Sympathetically mediated baroreflex branches were directly associated with their specific effector functions. Specifically, vBRS and mBRS positively correlated with average SVR and dP/dt_max_, respectively, while coBRS positively correlated with average cardiac output.

These findings suggest a functional compartmentalization of sympathetic activity. Supporting this, vasodilation in the contralateral forearm contrasts with increased vascular resistance in the calf during isometric exercise (Eklund et al., 1974; Rusch et al., 1981), reflecting separate central and peripheral mechanisms (Mark et al., 1985). Additionally, physical and mental stressors affect sympathetic nerve activity differently: isometric handgrip increases it in both muscle and skin. In contrast, mental tasks and cold pressure tests increase it only in muscle (McCarthy et al., 2024). In conditions like hypertension, obesity, and heart failure, sympathetic activity is elevated in muscles but not skin (Grassi et al., 1998; Grassi et al., 2009).

### 4.7. Positional Changes Influence on Baroreflex Branches

Transitioning from a supine to a standing position significantly reduced ibiBRS across delays from 0 to 6 beats after the SAP peak. This reduction was associated with increased heart rate, shorter LVET, and elevated DAP—mechanisms that counteract gravity and prevent orthostatic hypotension. These changes were accompanied by decreased RMSSD-HRV (indicating reduced parasympathetic activity) and increased LF/HF-HRV ratio (suggesting a more significant sympathetic influence). The highest baroreflex effectiveness in heart rate modulation occurred at the 0-beat delay in both supine and standing positions, underscoring the importance of rapid responses.

Postural changes diminished the baroreflex influence on the SA node, leading to parasympathetic withdrawal and increased heart rate to maintain blood pressure. Similar reductions in cardiovagal BRS have been observed during other orthostatic challenges (Dorantes-Mendez et al., 2013; O’Leary et al., 2003). Standing caused a modest decline in mBRS at 0-beat delay without any impact on dP/dt_max_, stroke volume, or cardiac output. However, this early parasympathetic activity may prepare the myocardium for a more robust sympathetic response as standing continues (Chapleau et al., 2011; Machhada et al., 2017).

Although BRS for muscle sympathetic nerve activity increases following head-up tilting (O’Leary et al., 2003), vBRS for SVR remained unchanged during standing, possibly due to undetectable transient vascular changes (Fu et al., 2006) or effects on venous volume rather than arterial resistance (Johnson et al., 1974; Lacolley et al., 1992). However, the peak vascular baroreflex effectiveness (vBEI) and frequency (vSQ) were delayed from 1-2 to 3-6 beats, suggesting an adaptation to altered hemodynamics. Overall, postural changes primarily affect parasympathetic baroreflex control of heart rate, with minimal impact on sympathetic myocardial and vascular branches.

### 4.8. Limitations

This study has several limitations. First, the relatively small sample size, particularly from the EUROBAVAR dataset, may limit the generalizability of the findings. To functionally validate BRS estimations, we used the preoperative dataset to assess BRS associations with cardiovascular variables and the EUROBAVAR dataset to evaluate the effects of orthostatic challenges. However, the heterogeneity and female majority in the EUROBAVAR group may affect the robustness of the results.

Additionally, while the noninvasive methodology provides valuable insights into baroreflex dynamics, it lacks the precision of invasive techniques, such as direct sympathetic nerve activity recordings; this may limit the ability to fully capture the nuances of autonomic regulation, particularly with sympathetic pathways.

Lastly, the cross-sectional design prevents the study from assessing longitudinal changes in baroreflex function, which would provide deeper insights into autonomic regulation over time. Future research with larger, more homogeneous samples and longitudinal designs is needed to validate these findings further.

### 4.9. Conclusion

Our noninvasive method effectively captures multiple baroreflex responses and their temporal dynamics, highlighting distinct autonomic mechanisms and the impact of postural changes. Parasympathetic ibiBRS was linked to RMSSD-HRV at a 0-beat delay, while sympathetic vBRS showed strong associations with SVR, cardiac output, and RMSSD-HRV, particularly at a 3-beat delay and was the only parameter associated with SAP at 1-beat delay. Sympathetic mBRS was linked solely to dP/dt_max_ at 1-beat delay, and coBRS correlated with cardiac output and SVR at 0- and 3-beat delays. Postural changes affected the parasympathetic ibiBRS and marginally sympathetic baroreflexes (mBRS and vBEI). Future research should validate these findings in larger samples, explore longitudinal changes, and investigate neurobiological mechanisms across different physiological and medical contexts.

## Author contributions

**Heberto Suarez-Roca:** writing - original draft, review and editing, conceptualization, data curation, formal analysis, visualization, funding acquisition, methodology, and supervision. **Negmeldeen Mamoun:** writing - review and editing, data curation, methodology, funding acquisition, and project administration. **Joseph P Mathew:** writing - review and editing, funding acquisition, supervision, and project administration. **Andrey V Bortsov:** writing - review and editing, formal analysis and statistics.

## Acknowledgments

This work was supported by the US National Institutes of Health (grant No. R21NS112912 to JPM) and institutional funding from the Department of Anesthesiology, Duke University Medical Center. The authors declare no conflicts of interest.

## Data Availability

The data supporting the findings of this study are available from the corresponding author upon reasonable request and in accordance with institutional policies.

